# Large contribution of repeats to genetic variation in a transmission cluster of *Mycobacterium tuberculosis*

**DOI:** 10.1101/2024.03.08.584093

**Authors:** Christoph Stritt, Michelle Reitsma, Galo Goig, Anna Dötsch, Sonia Borrell, Christian Beisel, Daniela Brites, Sebastien Gagneux

## Abstract

Repeats are the most diverse and dynamic, but also the least well understood component of microbial genomes. For all we know, repeat-associated mutations such as duplications, deletions, inversions, and gene conversion might be as common as point mutations, but because of short-read myopia and methodological bias they have received much less attention. Long-read sequencing opens the perspective of resolving repeats and systematically investigating the mutations they induce. For this study, we assembled the genomes of 16 closely related strains of the bacterial pathogen Mycobacterium tuberculosis from PacBio HiFi reads, with the aim of characterizing the full spectrum of DNA polymorphisms. We find that complete and accurate genomes can be assembled from HiFi reads, with read size being the main limitation in the presence of duplications. By combining a reference-free pangenome graph with extensive repeat annotation, we identified 110 variants, 58 of which can be assigned to repeat-associated mutational mechanisms such as strand slippage and homologous recombination. While recombination events are less frequent than point mutations, they can affect large regions and introduce multiple variants at once, as shown by three gene conversion events and a duplication of 7.3 kb that involve ppe18 and ppe57, two genes possibly involved in immune subversion. Our study shows that the contribution of repeat-associated mechanisms of mutation can be similar to that of point mutations at the microevolutionary scale of an outbreak. A large reservoir of unstudied genetic variation in this “monomorphic” bacterial pathogen awaits investigation.

## Introduction

DNA repeats are the most diverse and dynamic components of any genome, not counting viruses. They comprise a veritable zoo of elements of different origins and complexities, ranging from short tandem repeats and subtle palindromes to autonomously replicating transposable elements, inteins, and members of multigene families (for a comprehensive review see Treangen et al. 2009). While the molecular evolution of repeats is highly variable, they share the property of providing a substrate for homologous or illegitimate recombination. This makes them the principal cause of genome instability and DNA polymorphisms in the form of duplications, deletions, inversions, and non-reciprocal transfers between homologs through gene conversion (Darmon and Leach 2014).

Whole genome sequencing has shown that repeat-associated variation is ubiquitous. This insight stems from dedicated studies (e.g. Achaz 2002, Schmid et al. 2018), but maybe even more from frustrated attempts to assemble genomes or identify variants using reads that are too short to span repeats, which results in fragmented assemblies and ambiguous mappings. Along with short-read myopia comes a methodological bias towards point mutations, which are simpler to model and underlie downstream analyses such as phylogenetic inference and selection scans. For most organisms, little remains therefore known about the types, rates, and phenotypic effects of repeat-associated mutations. Recently, a systematic investigation of repeats has come within reach thanks to long-read sequencing and analytical tools such as pangenome graphs (Garrison, Guarracino, et al. 2023) and hierarchical alignment (Armstrong et al. 2020). For the streamlined genomes of prokaryotes, base-perfect assemblies can be created (Wick et al. 2023), and pangenome graphs can be used to obtain a concise representation of all variant types (Yang et al. 2023).

One area in which a full characterization of genetic diversity would be particularly useful is the study of bacterial pathogens. Some of the most deadly of them – including the agents of anthrax, typhoid, plague, leprosy, and tuberculosis – have been designated “monomorphic” because of their low levels of genetic diversity (Achtman 2012). Lack of horizontal gene transfer and evolution under extreme clonality contribute to the phenomenon of “monomorphy”, possibly through strong background selection (Stritt and Gagneux 2023). In the absence of horizontal gene transfer, intrachromosomal recombination between repeats might be a key mutational mechanism in these organisms. As in other organisms, however, the study of repeats and structural variants has been neglected since the advent of short-read sequencing, and we remain largely ignorant about their contribution to genetic and phenotypic variation.

In this study, we use Pacific Biosciences (PacBio) HiFi sequencing to characterize the full spectrum of DNA polymorphisms in 16 strains of *Mycobacterium tuberculosis*, the agent of tuberculosis (TB), which with other closely related lineages forms the *Mycobacterium tuberculosis* complex (MTBC, Gagneux 2018). We focus on a transmission cluster in the city of Bern, Switzerland, previously characterized through RFLP (Genewein et al. 1993) and Illumina sequencing (Stucki et al. 2015; Kühnert et al. 2018). The cluster reflects mainly transmission among homeless and substance abusers in the 1990ies, with spillovers to the general population and reactivated TB diagnosed up to 2012 (Stucki et al. 2015). This is not the typical manifestation of TB, which today mainly affects poor countries and is, after COVID-19, the second most deadly infectious disease in the world (WHO 2023). How these bacteria manage to be so successful with so little genetic diversity remains puzzling. Part of the answer may lie in the genomic “dark matter”, the repetitive 10% in the genomes of these bacteria.

Here we unlock the repeatome of the MTBC by addressing a simple question: what types of variants are there? More specifically, we 1) evaluate the accuracy of assemblies constructed from PacBio HiFi reads; 2) characterize the repeat landscape of the MTBC; and 3) describe the different types of DNA polymorphisms and their underlying mutational mechanisms. Our results show that the contribution of neglected genomic regions and types of mutations can be similar to that of point mutations in non-repetitive regions.

## Results

### Complete and accurate assemblies from CCS reads

16 strains from the Bernese outbreak, isolated between 1988 and 2005, were selected for sequencing on a PacBio Sequel II in circular consensus sequencing (CCS) mode (Supplemental Fig. S1, Supplemental Table S1). Using the Flye assembly algorithm, all but one genome (P001-N1377) could be assembled into single circular chromosomes, despite considerable variation in read lengths and sequencing depths (Table 1).

**Table 1:** Sequencing and assembly statistics.

| Strain | Reads |  |  | Assembly |  |  |
| --- | --- | --- | --- | --- | --- | --- |
|  | Nr Reads | N50 | Longest | Length | Coverage | Circular |
| P007-N1108 | 44,212 | 5,344 | 21,567 | 4,405,381 | 47 | Y |
| P028-N1362 | 43,856 | 5,961 | 27,471 | 4,405,379 | 56 | Y |
| P003-N1374 | 11,692 | 6,726 | 24,804 | 4,405,378 | 16 | Y |
| P001-N1377 | 26,848 | 6,839 | 27,922 | 4,412,646 | 38 | N |
| P008-N1380 | 24,578 | 7,534 | 27,155 | 4,405,372 | 38 | Y |
| P010-N1385 | 53,811 | 5,391 | 21,423 | 4,405,379 | 58 | Y |
| P020-N1386 | 29,983 | 7,884 | 27,587 | 4,405,380 | 48 | Y |
| P059-N1392 | 76,540 | 4,781 | 15,395 | 4,405,268 | 70 | Y |
| P073-N1394 | 38,388 | 7,422 | 27,900 | 4,405,232 | 59 | Y |
| P074-N1402 | 31,862 | 7,594 | 29,206 | 4,405,232 | 50 | Y |
| P066-N1411 | 26,719 | 7,075 | 28,501 | 4,405,380 | 39 | Y |
| P034-N1426 | 66,552 | 6,740 | 27,519 | 4,405,492 | 90 | Y |
| P052-N1429 | 26,244 | 7,941 | 27,868 | 4,405,378 | 43 | Y |
| P042-N1430 | 18,475 | 7,657 | 26,514 | 4,405,378 | 29 | Y |
| P022-N1431 | 40,133 | 7,474 | 27,860 | 4,405,379 | 62 | Y |
| P006-N1591 | 42,575 | 7,713 | 27,040 | 4,405,380 | 68 | Y |

To test whether assemblies are not only complete but also accurate, we aligned the long reads back against the respective assemblies and called variants to discover inconsistencies between the two. Five sites in four assemblies were identified where the reads contradict the assembly, while 12 assemblies are free of inconsistencies and appear to be correct to the base. A closer inspection of the inconsistencies suggests that they are due to different causes (Supplemental Fig. S2). In P003-N1374, the assembly with the lowest mean coverage (16x), a total of five reads disagree on a sequence of five versus four cytosines. A second case, in P028-N1362, seems to reflect genuine heterogeneity as it might arise during culture or from a heterogeneous inoculum, with 28 reads supporting an adenine versus 28 reads supporting a guanine. A third inconsistency arises from a duplication in P001-N1377 (discussed below). This sequence was not resolved by Flye but in the subsequent circularization step, where 12 bp evident in the reads went missing. Finally, two nearby single-base insertions in P034-N1426 suggested by the assembly but not the reads reflect misassembly: one single read shows the presence of the two additional bases, while 79 contradict it. The last two inconsistencies, which are clear assembly errors, were corrected: 12 bp were added to the duplication in P001-N1377, and two bases were deleted from P034-N1426.

### The repeat landscape of Mycobacterium tuberculosis

Considering the key role of repeats in causing structural variation and gene conversion, we first sought to understand the repeat landscape in the studied genomes. Different types of repeats were annotated in one newly assembled genome, P034-N1426 (Fig. 1): homopolymers of at least 5 bp, short sequence repeats (SSRs, direct repeats of 3 to 9 bp), tandem repeats (TRs, direct repeats > 9 bp), and insertion sequences. Homopolymers were the most abundant type with 6,770 occurrences. 67 SSRs were identified, the large majority of them triplets of six or nine that do not shift reading frames (Fig. 2A). TRs were found at 47 locations, with quite some variation in repeat periodicity and length (Fig. 2A). Finally, 57 insertion sequences were identified, including 12 copies of IS6110 (IS3 family) and 9 copies of IS1081 (IS256 family), two IS families in the MTBC that are known to vary in copy number.

**Figure 1:**
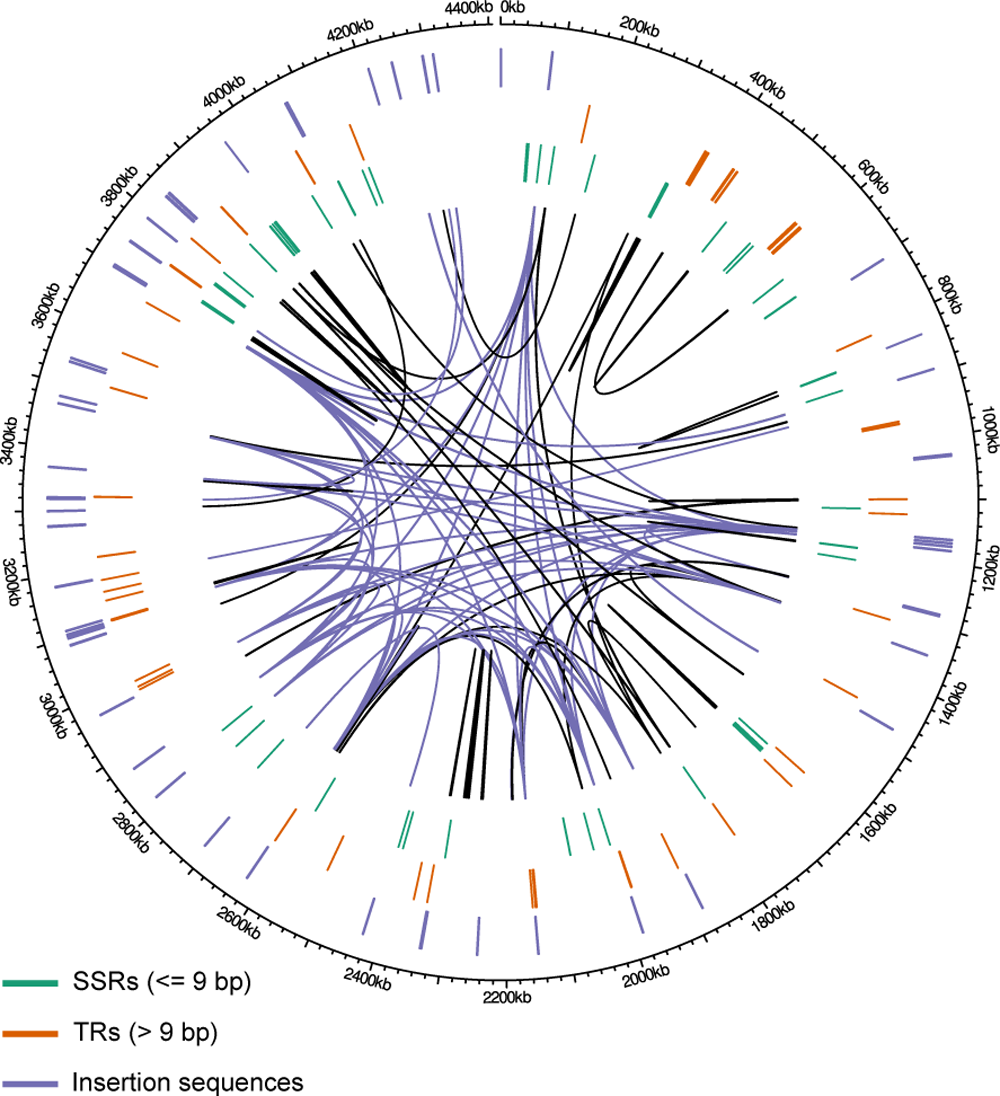
Repeat landscape in P034-N1426. Annotated short sequence repeats (SSRs, <=9 bp), tandem repeats (>9 bp), and insertion sequences are shown in the three outer tracks. Homopolymers are not shown because of their large number. The links inside the circle show pairs of homologous sequences of at least 90% identitiy and 50 bp length. Links that connect copies of insertion sequences are shown in purple, others in black.

**Figure 2:**
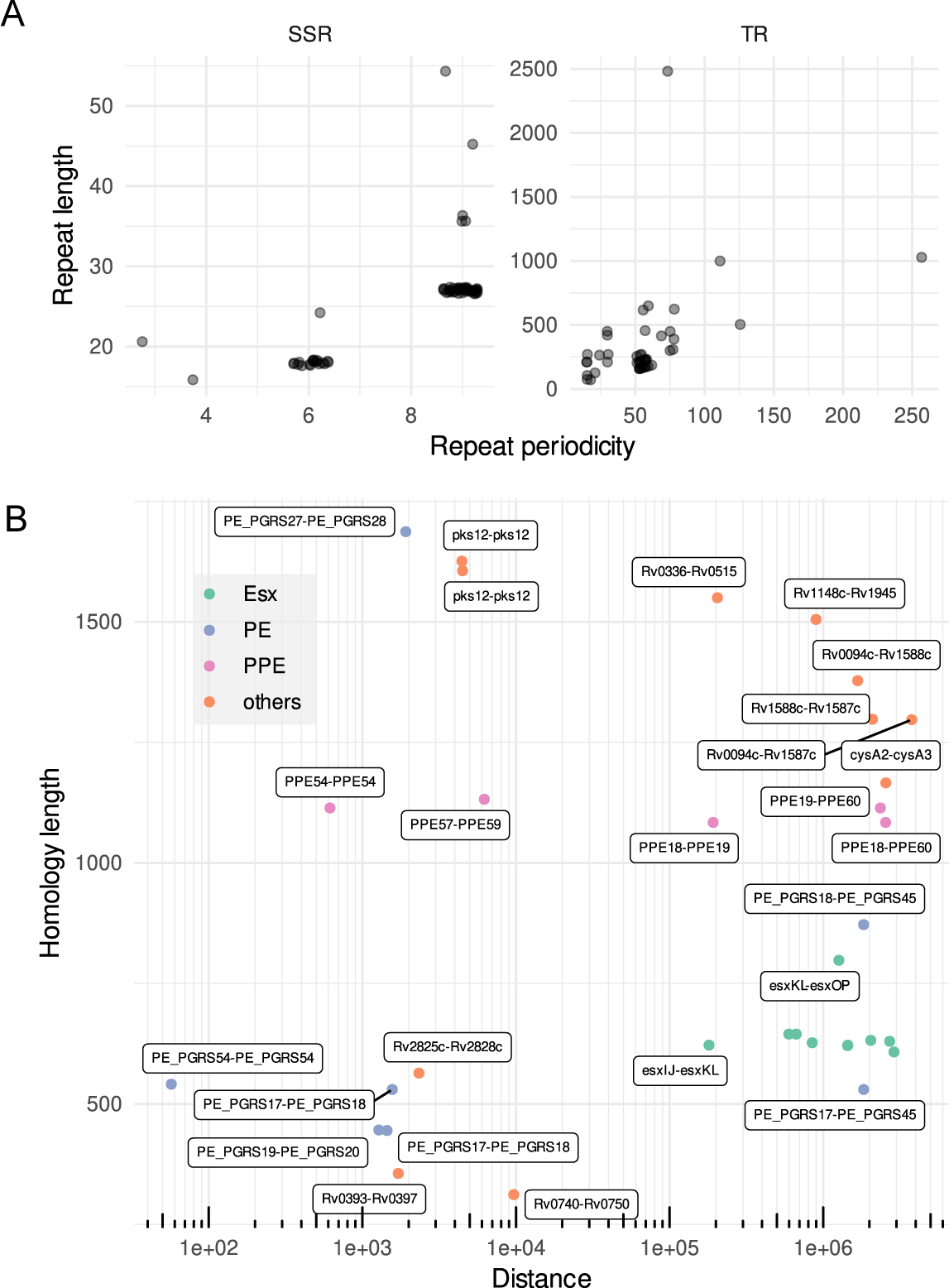
Characteristics of SSRs, TRs, and homology pairs. A) Periodicity and total length of short sequence repeats (SSRs) and tandem repeats (TRs). B) Gene pairs sharing substantial homology (over at least 20% of their length). The x-axis shows the distance between the pairs, the y-axis the length of the homology segment. Colors indicate the three main repetitive gene families in the MTBC. Not all *esx* pairs are labeled due to lack of space.

As a second approach to characterize the repeat landscape, we identified pairs of homologous sequences of at least 50 bp and 90% identity across the genome. Sequence homology is the substrate for homologous recombination and thus informative about where in the genome we might expect recombination-associated mutations. The thresholds are somewhat arbitrary as the minimal efficient processing segment (MEPS) for homologous recombination is not known in the MTBC; this point is addressed in the discussion. Excluding homology pairs within TRs to avoid redundancy in repeat annotation, we identified 136 pairs of homologous sequences, making up a total of 93,685 bp or 2.1% of the 4.4 Mb genome (Fig. 1).

To better understand which genetic elements share sequence homology, we intersected the homology segments with the gene and IS annotations. The repeat landscape is dominated by 45 and 15 homology pairs of highly similar IS6110 and IS1081 copies, respectively (Fig. 1). Members of the ESX, PE, and PPE gene families constitute a second prominent feature: 9 ESX genes (*esxI*, *esxJ*, *esxK*, *esxL*, *esxN*, *esxO*, *esxP*, *esxV*, *esxW*), 8 PPE genes (*ppe18*, *ppe19*, *ppe34*, *ppe38*, *ppe46*, *ppe57*, *ppe59*, *ppe60*), and 7 PE_PGRS genes (*pe_pgrs17*, *pe_pgrs18*, *pe_pgrs19*, *pe_pgrs20*, *pe_pgrs27*, *pe_pgrs28*, *pe_pgrs45*) are part of homology pairs (Fig. 2B). Most of the remaining sequence homology is found between pairs of genes of unknown function, designated by gene names beginning with “Rv” in Figure 2B.

### Types, frequencies, and genomic context of the 110 identified variants

Equipped with a basic understanding of what types of repeats are located where in the genome, we constructed a pangenome graph from the 16 assemblies and identified “bubbles” in the graph. 110 variants at 109 sites were identified (Fig. 3A), including 75 single nucleotide polymorphisms, 11 multinucleotide polymorphisms (MNPs, simultaneous changes of two or three base pairs), 17 deletions and 7 insertions. Deletion and insertion lengths range from 1 to 7,346 bp, the majority being indels smaller than 10 bp (Fig. 3B). Regarding the frequency of the variants, 88 of 110 are singletons, that is, are present in only one of the sampled strains, 14 are shared between up to five strains, and eight are present in all but P034-N1426, the strain that diverged early from the rest of the sample (Fig. 3C, Supplemental Fig. S1).

**Figure 3:**
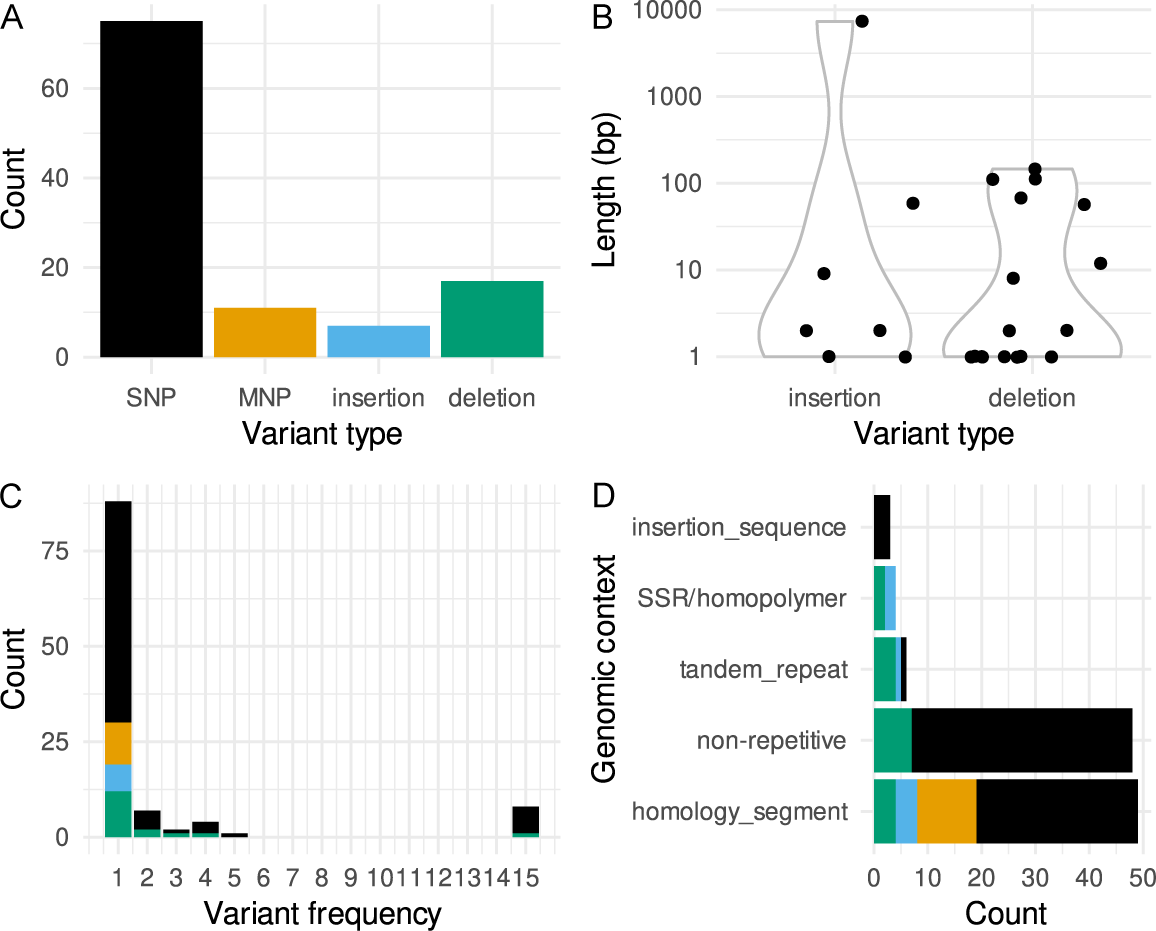
Types of DNA polymorphisms in the 16 sampled strains. A) Counts of SNPs, MNPs, insertions and deletions. B) Length of insertions and deletions, with a log-scale on the y axis. C) Frequency of the variants in the 16 strains. D) Intersection of the variant sites with the annotated repeats.

To identify repeat-associated mutations, we intersected the variant sites with our repeat annotation. 62 of the 110 variants are associated with repeats, while 48 occur in non-repetitive genic or intergenic regions (Fig. 3D). For most repeat-associated variants, the annotation directly suggests an underlying mechanism: of the seven insertions and deletions larger than 50 bp, five locate to tandem repeats, with the length of the variants corresponding to a multiple of the repeat periodicity. Four small indels are located in SSRs and homopolymers, suggesting strand slippage as underlying mechanism. The most striking pattern, however, is the large number of variants that intersect with homology segments. 49 variants (45%), including all MNPs, locate to homology segments. This is a more than 20-fold over-representation, considering that these segments make up 2.1% of the genome.

### Gene conversion between PE/PPE genes accounts for more than a third of all variants

A closer look at the variants occurring in homology segments shows that they occur as dense clusters of variants in single strains and are located in PE/PPE genes, two multigene families that are characteristic of pathogenic mycobacteria and play a role in host-pathogen interactions. 34 of the variants identified in P059-N1392 cluster in the repetitive C-terminal domain of *pe_pgrs28* (Fig. 4). Similarly, six variants in P003-N1374 cluster in *ppe18*, and four variants in P052-N1429 in *ppe19*. Since *ppe18* was annotated as a surface antigen, we further investigated the variants in this genes and found that two non-synonymous mutations affect two distinct epitope regions in the gene (Supplemental Fig. S3).

**Figure 4:**
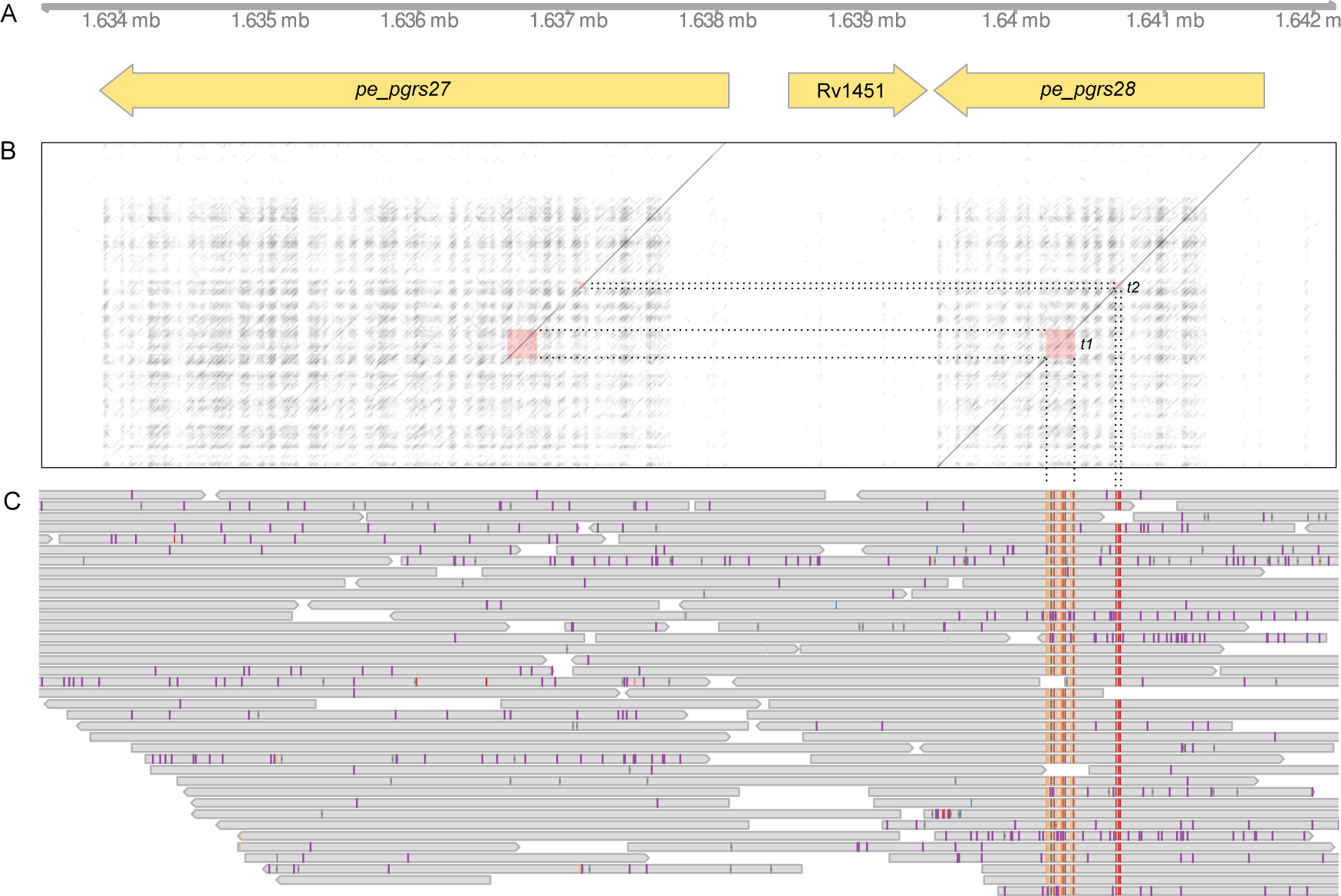
Gene conversion event between *pe_pgrs27* and *pe_pgrs28* in the strain P059-N1392. A) Genes annotated in the affected region. B) A sequence dotplot of the whole region in P059-N1392 on the x-axis and the sequence of the target gene *pe_pgrs28* on the y-axis. The red squares show the two conversion tracts t1 and t2. C) PacBio reads of P059-N1392 aligned against P034-N1426, highlighting the 34 variants in the two conversion tracts.

Given that these variant clusters occur in segments of homology, we hypothesized that they were caused by gene conversion between close paralogs. To test this, we blasted the suspected conversion tracts against the genome, expecting two exact hits in the two paralogs involved versus only a single exact hit in the source gene in strains where no gene conversion occurred. Indeed the conversion tract in *pe_pgrs28* yields two exact matches in *pe_pgrs27* (Fig. 4). The matching regions are separated by 286 bp, suggesting that the conversion tract is not continuous but interrupted by a stretch of the target gene. The suspected conversion tracts in *ppe18* and *ppe19* yield exact matches in *ppe19* and *ppe18*, respectively – gene conversion has worked both ways in different strains between these two genes, which are 190 kb apart.

### Birth of a new PPE gene through homologous recombination

As noted above, for one strain (P001-N1377) the Flye assembly step resulted in a single non-circularized contig. To understand why the assembly failed in this strain, we aligned the reads of P001-N1377 against a close relative where this region posed no problems. Double coverage and clipped reads, where the clipped parts feed back into the repeat on the opposite site, suggest that this region is duplicated in P001-N1377 (Supplemental Fig. S4). A comparatively long read would be required to resolve this 2 * 7, 346 14, 692 bp region, given an N50 read length of 6,839 bp for this strain. Indeed there is one read of 18,797 bp that spans the region (Supplemental Fig. S4). The short overlap on the 3’ side (140 bp) might explain why Flye failed to close the gap.

The duplication occurred in a region of the genome that contains multiple PE/PPE genes and insertion sequences (Fig. 5A) and where nested repeats testify to past duplication events (Fig. 5D). According to our de novo annotation, the duplication contains eight reading frames, five of them coding for PE/PPE genes. A comparison with the H37Rv reference annotation and our insertion sequence annotation shows that three CDS are part of a IS21 insertion sequence, while the unnamed gene in front of the IS is *ppe58* (Fig. 5A). Three of the five annotated PE/PPE CDS are not annotated in H37Rv; their sequences are similar to the closely related *ppe57*, *ppe58*, and *ppe59* (Fig. 4D), suggesting that these are leftovers from a previous duplication event.

**Figure 5:**
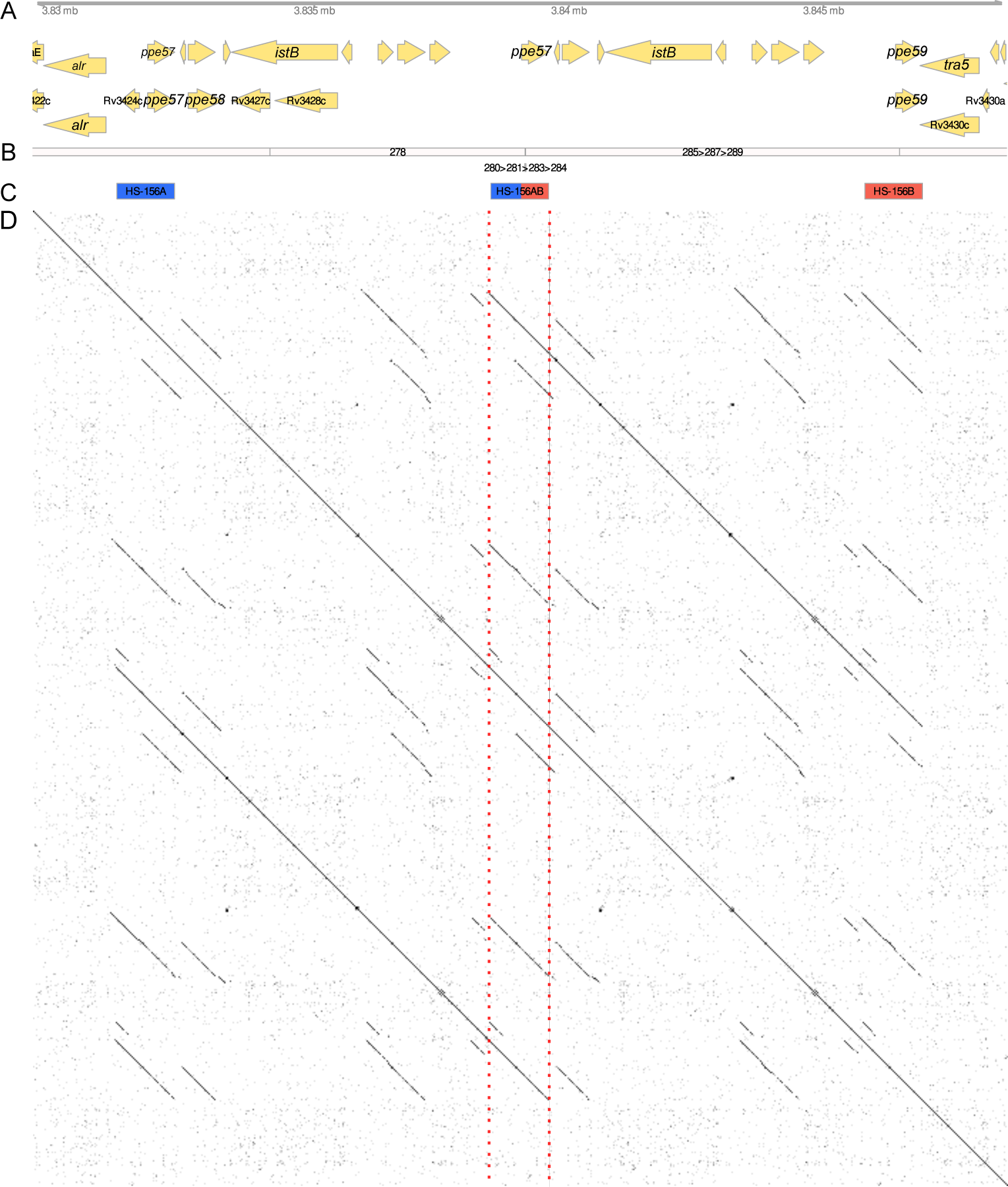
The duplicated region in P001-N1377. A) Genes annotated in the duplicated region, above from our *de novo* annotation, below from a liftover from the H37Rv reference annotation. B) Path in the genome graph corresponding to the duplicated region. C) Homology segments. D) Sequence dotplot of the duplicated region against itself. The red vertical lines highlight the center of the duplication where the duplicates overlap and the recombinant homology segment is located.

The gene models (Fig. 4A) proved not very helpful when trying to understand the convoluted graph for the duplicated region and how the duplication might have originated. More informative were the segments of homology (see above). A homology pair is present in the region, HS-156A and HS-156B (Fig. 5C); the two segments comprise *ppe57* and *ppe59*, respectively, and an additional stretch at the 5’ side of these genes. A blast search of these segments against the graph reveals a third segment at the very center of the duplication, highlighted by the red lines in Figure 4D, which is a recombinant between the two parental segments (Fig. 6A). The location of the recombination breakpoint is fuzzy but has to be located before or within the first 70 bp the new, identical copy of *ppe57*, as *ppe57* and *ppe59* are identical for the first 70 bp but then differ substantially. The presence of a third homology segment at the center of the duplication suggests a mechanism of duplication through homologous recombination (Fig. 6B).

**Figure 6:**
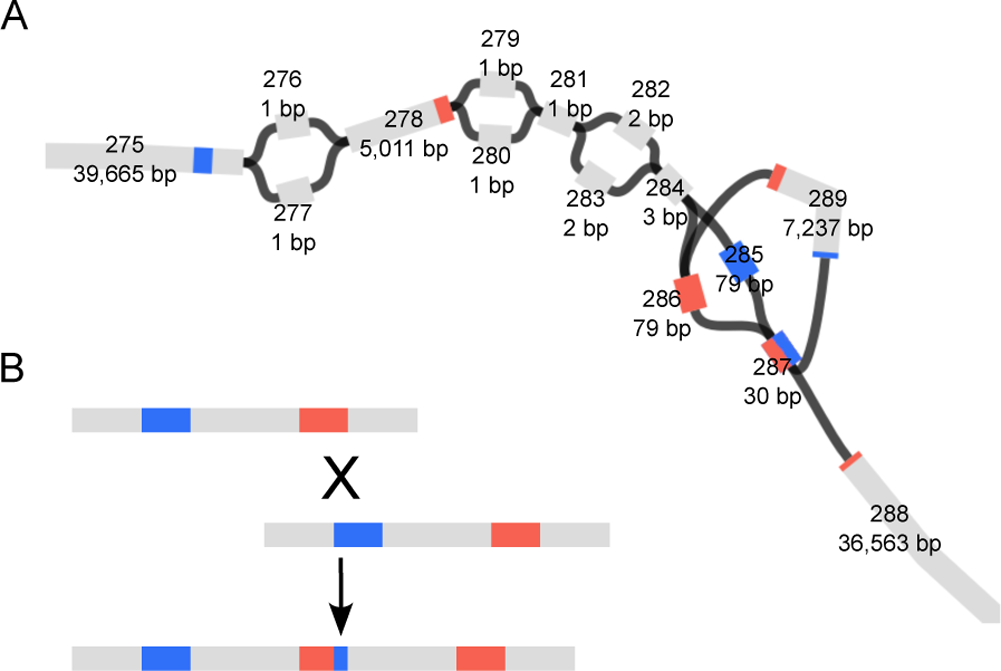
A) Graph representation of the duplication. Nodes are numbered and their length in bp is indicated. The colors indicate sequence similarity as determined through a blast search of the duplicated and adjacent parts. B) A model of homologous recombination underyling the duplication.

## Discussion

The promise of third generation sequencing technologies is to resolve repeats and thus enable a systematic study of repeat-associated genetic variation. In this study we show that virtually base-perfect MTBC genomes can be assembled from PacBio HiFi reads. These provide the basis to explore all genetic variation present in a set of genomes, in particular in the repeatome. In the following, we discuss homologous recombination as a mechanism underlying genetic diversity and argue that this mechanism is frequent and prone to have fitness consequences for the bacteria. We then situate our study among other recent attempts to obtain complete genomes from long reads and discuss read size as the limiting factor in the presence of duplications. Finally, we consider the use of complete genomes in genome-scale inference and point out caveats when including repeat-associated mutations for estimating phylogenies and selection scans.

### Recombination re-emerges as an important mechanism in the MTBC

Our analysis of a sample of 16 strains in a single transmission cluster does not lend itself to generalizations. Still, the observation of three gene conversion events, a duplication, and several insertions and deletions in tandem repeats in a sample of closely related strains suggests that homologous recombination operates frequently. While point mutations remain the most frequent events (Fig. 3A), recombination can affect large regions and contribute disproportionally to genetic variation: three gene conversion events between members of the PE/PPE families (*pe_pgrs27*/28, *ppe19* and *ppe18*) account for 44 of the 110 variants identified, while the largest variant is a duplication of 7.3 kb that arose from recombination between a homology pair encompassing *ppe57* and *ppe59*.

Evidence for gene conversion and homologous recombination was presented in a series of studies in the early 2000s, before the topic largely disappeared with the advent short read sequencing. Karboul et al. (2006) first discussed gene conversion as a potentially important mechanism in the MTBC and described recurrent conversion between the adjacent *pe_pgrs17* and *pe_pgrs18*. Evidence for gene conversion between members of the PPE (McEvoy, Van Helden, et al. 2009) and ESX (Uplekar et al. 2011) families followed, as well as for *pe_pgrs27* and *pe_pgrs28* (Delogu et al. 2008), the gene most strongly affected by gene conversion in our study, and *ppe18* and *ppe60* (McEvoy, Cloete, et al. 2012).

These and similar studies were based on PCR amplification of few genes. With short read sequencing, the study of repeats and homologous recombination was marginalized. Several studies still investigated “recombination”, but used this general term to denote horizontal gene transfer (HGT) rather than intrachromosomal recombination (e.g. Godfroid et al. 2018; Chiner-Oms et al. 2019). The apparent absence of HGT in the MTBC is expressed in the paradigm “TB does not recombine”. As repeats are now coming into focus again, it is important to recall that homologous recombination is a many-sided fundamental mechanism involved not only in HGT, but also in replication and repair, and its underlying pathways are present in the MTBC (Gupta et al. 2011).

Some expectations regarding the occurrence of repeat-associated mutations, and particularly homologous recombination, can be formed by considering the distribution of repeats in a single genome (Fig. 1). The rate of recombination, and thus gene conversion, decreases exponentially as sequences diverge (Shen and Huang 1986). Indeed, to our knowledge all examples of gene conversion in the MTBC involve closely related genes. Even in the large PE and PPE gene families, with 99 and 68 members in the reference strain H37Rv, close homology is restricted to relatively few pairs and triplets of genes, and to some larger repeats within single genes (*pe_pgrs24*; Fig. 2B). These numbers would be higher with more permissive thresholds for homology search; e.g. *ppe58* would appear as a paralog of *ppe57* and *ppe59* (Fig. 4 Fishbein et al. 2015).

While the number of genes involved in recombination might be small relative to the size of these gene familes, at least some of the variants in these genes will have phenotypic consequences and affect traits of primary interest in MTBC research, including antimicrobial resistance, immune response, and virulence. The three main repetitive gene families in the MTBC (PE, PPE, ESX) play important roles in host-pathogen interactions; they code for secretion systems, surface receptors or secreted proteins that interact with the human immune system and were instrumental in the evolutionary transition to an obligately pathogenic lifestyle (Gey Van Pittius et al. 2006). Even in our small sample of closely related strains we found two genes affected by gene conversion and duplication, *ppe18* and *ppe57*, for which there is experimental evidence for a role in immune subversion (Nair et al. 2009; Xu et al. 2015). Gene conversion has been shown to be a cause of antigenic variation in different prokaryote species (reviewed by Santoyo and Romero 2005). Given that both *ppe18* and *ppe57* are part of TB vaccines in active development (Guo et al. 2023), a better understanding of their molecular evolution could be of practical relevance.

One type of variant we did not observe are insertion sequence polymorphisms, even though copies of IS6110 and IS1081 are the most conspicuous features of the repeat landscape in the studied genomes. IS6110 is an active element that varies in copy number in the MTBC, from zero to more than 25 (McEvoy, Falmer, et al. 2007)). Copies of this element flank large inversions and deletions (Roychowdhury et al. 2015), suggesting that they are involved in homologous recombination. Because IS copies are dispersed through the genome, recombination between copies with other outcomes than gene conversion is likely to be disruptive and would be rarely observed. It will be interesting to test whether high copy numbers of IS6110 lead to more deletions and inversions or, on the opposite, high copy numbers are made possible by mutations that decrease the recombination rate and thus the rate of deleterious large-scale structural variants.

### Duplications are the new dark matter

Our results show that assemblies from PacBio CCS reads are complete and essentially error-free. One single of the 16 genomes could not be closed in the primary assembly step, and only five sites in a total of 70,493,034 assembled bases were found to be ambiguous or inconsistent with the underlying reads. Furthermore, surprisingly low sequencing depths are required to obtain complete MTBC genomes: a depth of 16x was sufficient to obtain a closed genome with only one single ambiguous position. These assemblies are thus an appropriate point of departure to study the molecular evolution of repeats and dynamic gene families.

The accuracy of CCS reads allowed us to forego many of the methodological complexities of previous approaches to long-read assembly. Noisy long reads necessitate high sequencing depths and hybrid approaches that combine long and short reads and multiple assembly and polishing steps (e.g. De Maio et al. 2019, Wick et al. 2023). This results in complicated pipelines and directs attention away from biological to technical questions. Our results show that a simple approach that combines CCS reads and the Flye assembly algorithm performs well, resulting in assemblies that can be trusted.

The principal limitation of CCS reads, at present, is their size and thus the ability to bridge duplications and amplifications (Tvedte et al. 2021). Duplications, thus, are the new dark matter and the reason why the assembly problem is not solved for the MTBC. The proportion of long reads might be increased through improved DNA extraction and library preparation (Wick et al. 2023). But even with longer reads, whether an assembly can be closed also depends on the frequency and length of duplications and amplifications.

In the strains analyzed in this study, we stumbled upon a duplication of 7,346 bp that was bridged by a single read. Compared to other duplications that have been described in the MTBC, this is not particularly large, but its assembly still requires reads longer than 14,692 bp. Well described duplications are the 30 kb DU1 and 36 kb DU2 duplications in the BCG vaccine clade (Brosch et al. 2000), or the massive duplication of 350 kb that includes the DosR regulon (Domenech et al. 2010) and has appeared repeatedly in lineages two and four (Weiner et al. 2012). More recent examples are a 38- to 60-fold amplification of *esxR*/*esxS* and flanking PE/PPE genes in H37Rv mutant strains where the ESX-3 excretion system had been deleted (Wang et al. 2022); and a 120 kb duplication that evolved twice independently during experimental evolution (Smith et al. 2022). These genomes would not have been completed with the approach presented here. The most extensive investigation of amplifications in the MTBC so far, based on short-read coverage, suggests that amplifications are frequent but restricted to few genomic regions: 590 amplifications were found in 1,000 diverse MTBC genomes, the large majority of them in 24 hotspots regions (Abrahams et al. 2022).

### Making use of the repeatome

Of more immediate practical concern than the fitness effects of repeat-associated mutations is the use of the repeatome in genome-scale inference. In an epidemiological context, a key consideration is how repeats can be included to increase the genetic resolution for the inference of transmission chains (e.g. Modlin et al. 2021, Marin et al. 2021). It is evident that using complete assemblies and all types of polymorphisms increases the available information; it is less obvious exactly how this information should be used. Much of phylogenetic inference and selection scans is based on substitution models that apply to sequentially introduced point mutations (Yang 2014). As shown above (Fig. 4), gene conversion can introduce many variants in a single event.

While a gene conversion event can be informative about tree topology when it is shared between strains, modeling it through a substitution model will result in exaggerated branch lengths and biased time estimates. For the same reason, variants due to gene conversion should be excluded in clustering analyses based on SNP thresholds. Multiple variant types and mutational models could be used simultaneously in a Bayesian setting, as has been shown with SNPs and short indels (Redelings and Suchard 2007).

Caveats also apply to inferring positive selection on genes affected by gene conversion. On the one hand, codon models as well as the popular *d_N_*/*d_S_* statistic are based on the same substitution models as phylogenetic inference. Applying these methods to close paralogs can reveal false signatures of positive selection (Casola and Hahn 2009), and we suspect that previous reports of positive selection acting on a large number of PE/PPE genes (e.g. Zhang et al. 2011 Namouchi et al. 2013, Phelan et al. 2016) might have been exaggerated. Also homoplasy, a second popular signature of selection, has to be interpreted with care since gene conversion mimics convergent evolution (Green et al. 2023).

## Methods

### Samples and genome sequencing

Strains for PacBio sequencing were selected to represent the “Bernese outbreak”, a small transmission cluster belonging to lineage 4 and including 68 patients in the city of Bern (Switzerland), sampled between 1988 and 2011 (Figure S1, Supplementary Table S1; Genewein et al. 1993, Stucki et al. 2015). We first estimated a phylogenetic tree using the previously published short reads (NCBI bioproject PRJEB5925). A SNP alignment was created as previously described (Gygli et al. 2021), ignoring variants in known resistance genes and repetitive regions. The resulting alignment of 142 variable positions was used to estimate a phylogeny with raxml-ng v. 1.2.1 (Kozlov et al. 2019), using the GTR substitution model and 100 bootstrap replicates. Branch lengths were rescaled to account for non-variable positions.

16 strains from the different subclusters were selected for sequencing. For two patients (P022, P028), different isolates than the ones used by Stucki et al. were accidentally selected. Since the aim of this study is not epidemiological inference, we still used them and unambiguously labelled the assemblies with the patient number and the isolate name separated by a dash.

For DNA extraction, strains were grown for two to three weeks on 7H11 plates inocculated from 7H9 liquid wake-up cultures. DNA was extracted from the plates using a CTAB method (Van Embden et al. 1993) that includes RNAse treatment and a purification step using magnetic beats. SMRTbell libraries were prepared at the Genomics Core Facility jointly ran by ETH Zurich and the University of Basel and sent for sequencing on a Pacific Biosciences Sequel II System at the Lausanne Genomic Technologies Facility.

### Genome assembly and variant calling

Read quality statistics were obtained with LongQC (Fukasawa et al. 2020). We used Flye v. 2.8.1 (Kolmogorov et al. 2019) to assemble genomes from circular consensus sequences (CCS) and reoriented the assemblies to start with *dnaA* using Circlator v. 1.5.5 (Hunt et al. 2015). As *M. tuberculosis* is not known to contain plasmids, assembly is expected to result in a single circular chromosome.

A pangenome graph representation of the assembled single-contig genomes was build using PanGenome Graph Builder (pggb) v. 077830d (Garrison, Guarracino, et al. 2023); for an evaluation of pggb in bacteria see Yang et al. 2023). Percentage identity (-p) was set to 99, segment length (-s) to 5k, according to the high similarity expected in strains from a single transmission cluster and the maximum length of repeats in the genome. An arbitrary strain, P034-N1426, was used as a positional reference for outputting variants with *vg deconstruct* and for the annotation of repeats (see below).

To classify structural variants as insertions or deletions, we estimated the ancestral states of the variants by comparint them to a closely related strain that was also sequenced for the present study: N1015, a strain from Nepal assigned to sublineage 4.4.1.2, while the Bernese strains belong to sublineage 4.4.1.1. The minor allele in the outbreak sample was assumed to be the derived state if it differed from the outgroup allele; if the minor equalled the outgroup allele, the major allele was assumed as derived.

### Assembly validation and curation

To evaluate the accuracy of our assemblies, we looked for inconsistencies between reads and assemblies by aligning the long reads back against the assemblies and calling variants. If the assembly is accurate, no variants should be found, while variants identified this way indicate errors during assembly, circularization, or the presence of true genetic variation in the culture. We used minimap2 2.24-r1122 (Li 2018) to align the reads and called variants with freebayes v. 1.3.4 (Garrison and Marth 2012), setting ploidy to 1. Read-assembly inconsistencies were scrutinized with the Integrative Genomics Viewer (IGV, Robinson et al. 2011) and those unequivocally identified as assembly errors were curated using the pysam library of Biopython. The curated assemblies where validated through a second round of variant calling.

### Gene and repeat annotation

Genes were annotated *de novo* in all assemblies with bakta v.1.8.2 (Schwengers et al. 2021). To compare gene models, we also lifted over the H37Rv reference annotation (ASM19595v2) to P034-N1426 with liftoff v1.6.3 (Shumate and Salzberg 2021). To further characterize the repeat context of the variants, we annotated different types of repeats in the positional reference strain P034-N1426: insertion sequences with ISEScan v.1.7.2.3 (Xie and Tang 2017), short sequence repeats (<= 9 bp) with kmer-ssr v. 0.8 (Pickett et al. 2017), tandem repeats (> 9 bp) with SPADE v. 1.0.0 (Mori et al. 2019), and homopolymers of at least 5 bp using our own script (see GitLab link above). To invesigate these repeats in other assemblies, annotations were lifted over using *odgi position* on the pangenome graph. The variant positions resulting from *vg desconstruct* and the different annotations were intersected with bedtools v2.30.0 (Quinlan and Hall 2010).

To identify pairs of sequence homology across the genome, we used *nucmer* (–maxmatch –nosimplify) and *show-coords* from the mummer4 tool (Marçais et al. 2018) and removed self- and overlapping hits. We further filtered out pairs with less than 90% sequence identity and an alignment length smaller than 50 bp, as well as pairs that overlapped with annotated tandem repeats. The locations of homology segments were intersected with the *de novo* gene and IS annotations, using bedtools v2.30.0 (Quinlan and Hall 2010), to identify genetic elements potentially involved in recombination.

### Identification of gene conversion tracts

To test whether a cluster of variants identified in the same strain is due to gene conversion, we extracted the suspected conversion tracts from the respective genomes using odgi v0.8.3-57-gfbdb4d23 (Garrison, Guarracino, et al. 2023) and samtools v.1.18 (Danecek et al. 2021). We then used blastn v. 2.12.0 (Camacho et al. 2009) to align the sequences against a) the genome in which the cluster was detected, b) a genome in the ancestral state. In case of gene conversion, we expect two matches in a, in the source and the target gene, and a single match in b, i.e. only in the source gene. Visualization of the gene conversion and duplication events was done using the R package gviz (Hahne and Ivanek 2016) and dotter (Sonnhammer and R Durbin 1995).

Epitopes in PPE18 were downloaded from the Immune Epitope Database (iedb.org, accessed 24.1.2024). Miniprot v. (Li 2023) was used to infer the location of the epitope peptides within the nucleotide sequences of the genes.

## Data access

Raw reads and assembled genomes were deposited on the European Nucleotide Archive (PRJEB73759). The assembly pipeline is available as a Snakemake workflow on http://git.scicore.unibas.ch/TBRU/PacbioSnake.

## Acknowledgments

We wish to thank the members of the Gagneux group for the helpful discussions and the team of the sciCORE Center for Scientific Computing at the University of Basel for access to their computing cluster. This work was funded through grants from the European Research Council, grant number 883582, and the Swiss National Science Foundation, grant numbers 310030_188888 and CRSII5_177163.

